# Institutionalizing an Evidence-Based Practice in a Selected Government Agency in Nigeria

**DOI:** 10.1101/2020.12.21.423750

**Authors:** E. O. Nwaichi, I. Chidebe, J. O. Osuoha, E. Chimezie, C. Mbaoji, C. C. Ifeanyi-Obi, E.O. Ugwoha, B. Odogwu, U. Okengwu, V. Olua, L U. Oghenekaro, O. E. Agbagwa, N. Frank-Peterside, M. A. Ozah, C. Raphael, I. Ugiomoh, P. A. Nwoha, B. A. Akpoji

## Abstract

The need to tackle the discrepancy between research outputs and the execution of the findings into real practice is a crucial factor in establishing evidence-based practice in a selected government agency. There is a need to increase the number of experts in our education sector who can synthesise, translate, and package the evidence for ready use by decision-makers and to foster entrepreneurship in Africa. To achieve this, the activities of a 28-man Evidence Leaders in Africa (ELA) team were recruited to drive the project through capacity building training and workshop sessions. Thereafter, a non-random purposive sampling technique targeted at policy makers at various government and non-governmental organizations was adopted as the study design. In all, purposive administration of 424 copies of questionnaire to individuals in different governmental and non-governmental organizations was done by the recruited personnel. The retrieved data from the questionnaire were analyzed using standard statistical method. Socio-demographic characteristics of respondents reveal 34% representation of (31-40 years) age bracket, with (57%) of them working in the University spanning over 2 to 5years (47%) work experience. Approximately 64% of the respondents are not aware of EIDM while 71% had no understanding of EIDM. As regards knowledge of EIDM, 36% had no knowledge of EIDM, 29% had moderate knowledge, 34% were beginners while approximately 1% had advanced knowledge of EIDM. Also, approximately 99% of the respondents have neither been trained nor involved in training others in EIDM. It was also observed that 8.5% of the respondents were policy-makers in the organization, yet 90.7% of the respondents agree that EIDM is useful in policy-making while 81.3% of the respondents engage their colleagues in EIDM. From this study, it is safe to infer that institutionalizing EIDM in NNMDA has numerous benefits as shown by the findings of this study. This will only be possible when all parties involved in producing and using research evidence are well informed and knowledgeable in EIDM.

## 1.0 Introduction

In the past two decades, most nations of the world, which includes considerable number of developing countries, are increasingly recognizing and embracing the importance and need to use rational and more rigorous processes to ensure that healthcare recommendations are informed by the best available research evidence (González-Block, 2004; WHO, 2007). The idea of using an evidence to inform policy is not new. What is new and interesting however, is the increasing emphasis that been laid on the concept over the years. Evidence based policy/evidenced-based practice ideally, is committed to replacing the old ideologically driven politics with rational decision making. Evidenced informed policy-making (EIPM) is an “Approach that aims to ensure that decision-making is well-informed by the best available evidence. It is characterized by systematic and transparent access to, and appraisal of, evidence as an input into policymaking”. (Lavis, *et al*., 2009). In other words, “Evidence-informed policy provides an effective mechanism to establish in a scientifically valid way, what works or does not work, and for whom it works or does not work”. (Sutcliffe & Court, 2005). Creating a positive and supportive policy environment has been instrumental in expanding access and implementation of successful programmes and outcomes.

It has been proved with convincing information that evidence from research can enhance health policy development by identifying new issues for the policy agenda, informing decisions about policy content and direction or by evaluating the impact of policy (Innvær *et al*., 2002; Hanney *et al*., 2003; Dobrow *et al*., 2004; Campbell *et al*., 2009). Irresistibly without doubt, its’ been noted that better utilization of evidence in policy and practice can help save lives through more effective policies that respond to scientific and technological advances, reduce poverty and improve development performance in developing countries. (WHO, 2004).

Nigeria government in line with global development and practices, has recognized the relevance of evidence-based policy as a critical requirement for the improvement of her agencies especially the country’s healthcare systems (FMOH, 2006; Uneke *et al*., 2009). The Nigeria Evidence-based Health System Initiative (NEHSI) was established as part of the government’s effort and support to actualize the comprehensive health sector reform. This initiative held series of meetings in 2004 in Nigeria by the International Development and Research Centre (IDRC) at both Federal, State and Local government area levels. The meetings revealed a pressing need, as well as strong local and development partner support, for the building of a responsive evidence-based health system, with emphasis on primary healthcare (NEHSI, 2007, 2009). The importance of NEHSI initiative to developing country like Nigeria cannot be overemphasized, NEHSI in Nigeria was a six-year initiative programme which began in 2008, it was developed to support a fair, effective and efficient primary healthcare system. It was designed to be implemented in the 36 states including the Federal Capital Territory (FCT), but it only took place in Bauchi and Cross River states (NEHSI, 2007)

It is common practice and believed that government make and implement policy, this status quo needs to change within the government system. The government and policy-makers need to understand the value of evidence, become more informed as to what evidence is available, know how to gain access to it and critically be able to appraise it (Davies, 2004). The process of incorporating evidence into policy-making is neither a simple nor a straight forward act. The process of integrating evidence into policy decision making is a complex, competitive and is frequently being influenced by many actors in which evidence plays just of many roles. In Nigeria context, the complexity is further deepened in the sense that there is ‘no clear definition’ of who a policy-maker is, the constraint that exist between a policy maker, institution/organization and individual in the use of research evidence for policy-making.

This is the reason why it is necessary to create awareness and institutionalize the use of evidence to policy-making among the policy-makers in Nigeria and initiate the mechanism that will stimulate their interest in the subject. The major constraint to use of evidence in policy and practice in government selected agency (health-related agency) in Nigeria is grossly deficient in the capacity development mechanism at organizational and individual levels, particularly the lack of formally trained human resources in the area of health policy among public health policy makers and health service managers (Uneke *et al*., 2009). This barrier is caused by the gap differences existing between researchers and policy makers, so an opportunity should be sought to bring researchers, policy makers and policy managers together in common forum such as discussion forum or joint training to consider issues concerning evidence research to policy and practice interface. The policy-makers should be trained to equip them better with necessary skills that will enable to effectively acquire and assess evidence necessary for policy-making.

Weak capacity in seeking, appraising and applying evidence remains one of the major barriers to research use in policy-making and programming in the health sector in Nigeria (EIDM Guidelines Kenya, 2016). In government Agency, research evidence can only be beneficial to the human populace when they are applied. Although the barriers to use of evidence are enormous, it’s been established that access barriers, institutional/organizational barriers and individual barriers amongst others affect the use of evidence to policy and practice. These include lack of translational research or evidence, moderate access to internet, journals and other repositories, no research fund or grant as government hardly make provision. According to selected studies done on the major institutional barriers to evidenced-based practice (EBP) includes workload (manpower), lack of resources (funds) and institutional financial support, other staff/management not supportive of EBP, lack of authority to change practice, lack of incentive to promote research, weak collaboration, and a workplace resistant to change etc. (Sapkato, 2014). On the individual constraint to barriers to use of evidence to policy and practice constitute lack of capacity building, insufficient research skills, not being conversant with modern trend in research methodologies, lack of enabling environment for career development and training and retraining of research officers to implement EBP. The unavailability or inadequacy of these sources of research and research incentives to acquire and assess research at institutional or individual levels constitute a major impediment in health policy and system research evidence use in policy-making (Uneke *et al*, 2010).

Within the last decade the World Health Organization (WHO) and many other agencies have been vigorously supporting the process of contextualizing evidence and translating it into policy in many developing countries including Nigeria (WHO, 2003; AHPSR, 2007). The Health Policy and System Research (HPSR) championed by WHO aims to produce reliable and rigorous evidence that helps to inform the many and varied critical decisions that must be made by policy makers on how to organize the health system and effect changes.

In Nigeria, HPSR concept is new and most health policy makers, health service managers, health researchers, and major stakeholders at government and non-governmental levels are yet to completely appreciate its value in policy making and practice. Individual and organizational levels constraints enumerated earlier are perceived to be major impediments in HPSR evidence use in the health policy-making process in Nigeria (Uneke *et al*, 2009). This seems to be a general problem in most developing countries (González-Block, 2004).

There is a general consensus that despite the challenges, evidence thrives in positive development. Evidence help to understand policy environment and how it changes. Evidence is known to produce better deliverable outcomes, so institutionalizing evidence-based practice in government selected institution will produce better outcomes. The aim and objective of this prioritized the use of valid, quality and credible evidence-based practice in government selected agency and promote the use of organizational incentives in HPSR evidence in policy making.

The Evidence Leaders in Africa (ELA) project is aimed at creating awareness, and promoting the use of evidence in policy formulation and implementation by African Governments. Evidence refers to It is considered that addressing the critical gaps existing between the holders of scientific/technological knowledge and the Government/policy makers are key to driving effective and solution-driven policies. Policy making is the process by which governments translate their political vision into programmes and actions to deliver outcomes (Northern Ireland, 2016). However, we need to critically examine how the scientific and technical knowledge generated so far from the multi-disciplinary research outputs has been linked to policy making in Nigeria. Truly, the science and technology expertise should play key roles in government policy formulation processes for improved decision outcomes.

The focus of Science and Technology (S & T) is to continually contribute to knowledge base via research output, providing evidence for both the scientific and non-scientific ecosystems. Concomitantly, evidence from S & T should be harnessed through interactions between researchers and policy makers to create strong linkages, and provide support on advocacy matters.

Evidence Informed Decision Making (EIDM) in Policy making results in evidence based policy recommendations, and is guaranteed to take science and technology to the next level. Conversely, the existing divide between researchers and policy makers, the lacuna caused by fluidity in responsibility by both actors, and the resultant lack of co-ordination in collective efforts has limited progress. emphasizes the urgency to bridge this gap. Clearly, the need for synchrony cannot be over-emphasized.

It is in the light of this that the African Institute for Policy Development (AFIDEP) and the African Academy of Sciences (AAS) is implementing ELA project, under the auspices of the William and Flora Hewlett Foundation to build capacity in EIDM across governments in East and West Africa. It is in view of the current scenario that the Nigeria Natural Medicine Development Agency signed up as the first Government Institution in West Africa to institutionalize the EIDM in policymaking with a well-developed and publicized Guidelines for Evidence Use. The essence is to apply research to policy by engaging policymakers and researchers in ways that will facilitate S & T to influence policy, and *vice versa*. It is our goal that this will build, and remain pivotal in consolidating a sustained progress in developing a new cadre of professionals who will learn to interface between law, legislature, advocacy, science, science communications, technology, agenda setting and policy formulation in Nigeria.

To this end, this project seeks to strengthen the technical capacity of policymakers in a selected government agency in identifying, accessing, evaluating, interpreting, synthesizing, and deploying evidence in decision-making and policy implementation while building the capacity of participating researchers and consolidating a relationship thereof.

## 2.0 Methodology

This project commenced with a call for interest to pull researchers with burning interest in development and evidence together. The leadership of identified government agency was engaged to secure commitment and interest to participate. These activities birthed a 28-man ELA team commissioned to participate in capacity building training and workshop sessions for the duration of the project with a commitment to apply learnt skills at work to deploy evidence in policymaking and implementation as well as think reliable evidence creation during the course of experimental/ research design. A Guidelines Drafting Committee (GDC) was commissioned in the second quarter of the project life cycle to use background documents, other national and international policy documents to develop the Draft Guidelines for Evidence Use. The GDC utilized skills learnt in the project to put together a robust Guidelines. Draft Guidelines was finalized by involving wide-stakeholder consultations having key actors at across Nigeria and beyond. It is important to state that this study was reviewed and approved by University of Port Harcourt Ethics Committee prior to start.

### 2.1 Sample Size and Sampling Techniques

#### 2.1.1 Sample Size

Sample size is a representation of a research population; which is a recognizable group or combination of elements (e.g. people, organizations etc.) that are of interest to the scholar (Hair *et al*., 2000). The target audience depict the research population from which the sample size will be drawn. However, the aim and purpose of the study to a large extent determines the sampling method. These study targets policy makers at various government and non-governmental organizations (this however serves as our research population) as such a non-random purposive sampling method has to be adopted.

This study sees to a purposive administration of 424 copies of questionnaire to individuals in different governmental and non-governmental organizations.

#### 2.1.2 Sampling Technique

Sample size (424) calculated using Cochran’s (1963) formula was conveniently drawn from our population of interest. However, all elements that constituted the sample size were heads and policy makers who were available at the various governmental and nongovernmental organizations. This hence led to the deployment of a non-probability sampling method referred to as convenience sampling also (Ezejelue *et al*., 2008).

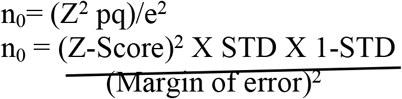

This equation is for an unknown population size or a very large population size N. Where the sample size(n_0_) is determined, assuming a 95% confidence level, 0.5 standard deviation, and a margin of error (confidence interval) of ±5% (0.05).

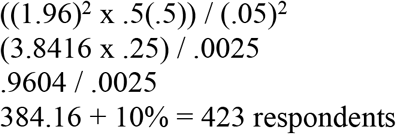

#### 2.1.2 Data Collection/Instrument Design

The instrument used for data collection is the questionnaire. A self-administered, questionnaire structured into sections A to F was used for this survey. Section A dealt with the Introduction of the researcher and ethical statement, section B – Socio-demographics of the respondents, while section C covered the study variables that has to do with the Knowledge and Awareness on Evidence Informed Decision Making (EIDM). Section D focused on variables relating to evidencebased practices, section E dealt with the study variables that pertains to perception of EIDM to policymakers, while section F covered the study variables relating to institutionalizing EIDM. The questions were in simple format thereby easing administration as they were also structured using five-point Likert scale which solicited information from the respondents capturing dependent, independent and moderating variables. Respondents consented to participation by agreeing to complete structured surveys as declared in the instrument. No minors were engaged in the study. The captured data from the questionnaire was analyzed using frequency and percentages, while mean was used to determine the mean response of the respondents for each of the questionnaire items. A five point Likert – type scale of Strongly Agreed=5, Agree = 4, Undecided = 3 Disagree =2, Strongly Disagree =1 etc., was used as rating scale. Response of Five point Likert-scale was categorized according to their mean score using the methodology of Agwu and Adeniran (2009) in terms of reliability.

The design of the study as shown in Figure 1 occurred in three main phases.

**Figure 1:**
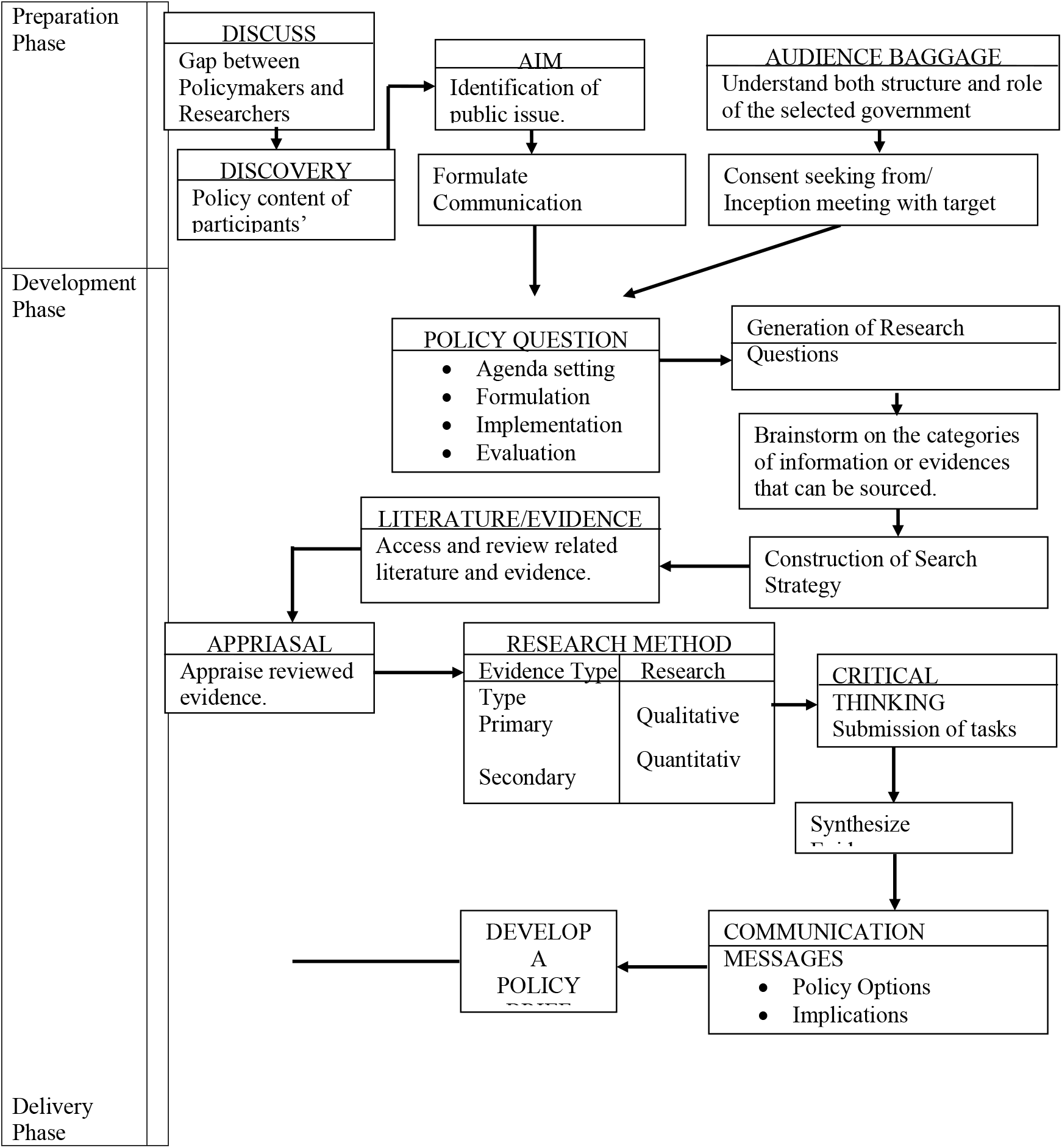
Study Design of Institutionalizing Evidence–Informed Practice

The preparation phase catered for the all preliminary activities that helped to chart the course of evidence-informed practice. The activities in this phase include:

i. Initiating a discussion among participants to give their individual perceptive on the gap that exists between policymakers and researchers. Participants were further challenged to identify the policy content of their individual research.
ii. The activity further led participants to point out the public issue to be addressed by evidence-informed research, which was a fundamental step that birthed the aim of individual works. The communication objectives that helped to achieve the aim were further formulated.
iii. Identification and understanding of target audience was a key activity in the preparation phase. It involved identifying the individuals and bodies who could bring the needed actions to reality, and they include a wide variety of persons such as, government officials, political leaders, news media, business leaders, community groups, amongst others. The target audience was further divided into primary audience, who directly affect the concerned policy, and secondary audience, who had influence on the primary audience. Having achieved audience baggage, strategic means to seek the necessary consent was considered; some means of communication included, calls, one-on-one appointment, echannels or text messages.

Activities of the preparation stage ushered in the development stage where the following activities took place:

iv. The policy question was developed by considering the different stages of the policy process such as; agenda setting where the magnitude of the problem and prioritizing of issues are made; formulation where the policy options are considered and a preferred option is selected; implementation, practical actualization of activities is reviewed on this stage and finally, monitoring and assessing of impact of an intervention is done in the evaluation stage.
v. Given that the policy question is broad in nature, the capacity of participants is built to be able to generate research questions that are specific to the policy question. The research questions informed the decision on where evidences can be sourced, which spanned over a wide range of sources, such as; media reports, academic papers, expert advice, official statistics, traditional knowledge, pilot studies and case studies.
vi. Related literature over a wide range of sources are hence reviewed using strategic search techniques such as converting concepts into search terms and using the Boolean search approach.
vii. Participants learn how to appraise evidence gathered in literature, the characteristics of individual designs and methods, as regards to its evidence and research type, are described and considered in appraising the strength of any sourced evidence. Critical thinking was taught to and applied by participants in appraising evidence, which further led to submission of tasks and activities by participants.
viii. Appraised evidences are represented in new ways that portray strong relationships and ease of interpretation. This representation was done by participants in evidence synthesis exercise. From the synthesized evidences, communication messages which include; policy options, implications and SMART recommendations are drawn, which were further used to develop the policy brief.

With the development of the policy brief, progress was made to the delivery phase.

ix. Strategic means of communicating the developed policy brief to the policymakers are explored such as power point presentations. Once participants have decided on which means of communication to be adopted, the actual presentation is done at the point of the policy window, where the presented policy issues are decided upon and acted on by the target audience.
x. Policy issues are continuously evaluated and monitored by participants.

#### 2.1.3 Data Collection/Instrument Design

The instrument used for data collection was the questionnaire. A self-administered, questionnaire structured into sections A to F was used for this survey. Section A dealt with the Introduction of the researcher, section B – Socio-demographics of the respondents, while section C covered the study variables that has to do with the Knowledge and Awareness on Evidence Informed Decision Making (EIDM). Section D focused on variables relating to evidence-based practices, section E dealt with the study variables that pertains to perception of EIDM to policymakers, while section F covered the study variables relating to institutionalizing EIDM. The questions were in simple format thereby easing administration as they were also structured using five-point Likert scale which solicited information from the respondents capturing dependent, independent and moderating variables.

#### 2.1.4 Data analysis

The captured data from the questionnaire was analyzed using frequency and percentages, while mean was used to determine the mean response of the respondents for each of the questionnaire items. A five point Likert – type scale of Strongly Agreed=5, Agree = 4, Undecided = 3 Disagree =2, Strongly Disagree =1 etc. was used as rating scale. Response of Five point Likert-scale was categorized according to their mean score using the methodology of Agwu and Adeniran (2009) in terms of reliability.

## 3.0 Results and Discussion

### 3.1 Socio demographic characteristics of respondents

Table 1 showed respondent’s demographic characteristics. The respondents were spread across almost all age brackets indicated with relatively large proportion (34%) within the age brackets of 31-40years. This is an indication that they are still at their productive stage of their career and need to be properly. The respondents were drawn from different workplace, mainly financial institutions (5%), clergy (2%), entrepreneurs (4%), health workers (9%), oil and gas (2%) and University (57%). As regards work experience, majority of the respondents had work experience of between 2 to 5years (47%) and 1year and below (42%). This implies that most of them are relatively new in their current position. Approximately 94% are permanent staff in their establishment with 99.1% having at least a first degree. This shows that the majority of the respondent were well educated. This could be an advantage in respondent’s interest and understanding of the importance of institutionalizing evidence-informed practice in Nigeria. Ashraf, Khan, Ali and Iftikham, (2015) noted that as the educational level of people increases, the interest to gather information also increases. Educated persons according to the author are able to take risks in businesses as well as easily adopt innovative techniques and practices.

**Table 1.** Respondents Socio demographic characteristics

| **Variable** | **Sub Variable** | **Frequency** | **Percentage** |
| --- | --- | --- | --- |
| **Age** | 21 – 30 yrs. | 83 | 24.5 |
|  | 31 – 40 yrs. | 116 | 34.1 |
|  | 41 – 50 yrs. | 85 | 25.0 |
|  | Above 50 yrs. | 56 | 16.5 |
| **Workplace** | Financial institution | 17 | 5.0 |
|  | Clergy | 8 | 2.4 |
|  | Entrepreneur | 14 | 4.1 |
|  | Government agency or ministry | 66 | 19.4 |
|  | Health workers (lab scientists, pharmacists, public health workers, laboratory scientists etc.) | 31 | 9.1 |
|  | Judiciary | 2 | 0.6 |
|  | NCDC | 1 | 0.3 |
|  | Oil and gas | 7 | 2.1 |
|  | Private institution | 2 | 0.6 |
|  | Research laboratory | 1 | 0.3 |
|  | University | 191 | 56.2 |
| **Duration of current position (years)** | 0 – 1 | 144 | 42.4 |
|  | 2 – 5 | 160 | 47.0 |
|  | 6 – 10 | 35 | 10.3 |
|  | 11 - 14 | 1 | 0.3 |
| **Job Category** | Admin staff | 25 | 7.4 |
|  | CEO | 14 | 4.1 |
|  | Clergy | 2 | 0.6 |
|  | Demography and area making, prevention of infectious disease | 1 | 0.3 |
|  | Field staff (extension services) | 1 | 0.3 |
|  | Front desk | 1 | 0.3 |
|  | Health and safety | 2 | 0.6 |
|  | Health worker | 11 | 3.2 |
|  | HR | 5 | 1.5 |
|  | Judge | 2 | 0.6 |
|  | Manager | 6 | 1.8 |
|  | Marketer | 5 | 1.5 |
|  | Marketing manager | 1 | 0.3 |
|  | Pharmacist | 11 | 3.2 |
|  | Policy maker | 38 | 11.2 |
|  | Researcher | 175 | 51.5 |
|  | Resident pastor | 4 | 1.2 |
|  | Rev. Minister | 1 | 0.3 |
|  | Snr Minister | 1 | 0.3 |
|  | Teacher | 1 | 0.3 |
|  | Technical staff | 22 | 6.5 |
|  | Technologist | 11 | 3.2 |
|  | The above section of the table is not necessary. If at all we needed this, we would have checked their cadre to know whether they are senior staff, middle cadre or junior staff. It gives us a view of whether our respondents by virtue of their cadre are involved in policy making |  |  |
| **Designation** | Permanent staff | 319 | 93.8 |
|  | Technical staff | 4.0 | 1.2 |
|  | Temporary staff | 6.0 | 1.8 |
|  | Volunteer staff | 11 | 3.2 |
| **Highest Educational Qualification** | Degree | 184 | 54.1 |
|  | Diploma | 3 | 0.9 |
|  | Masters | 86 | 25.3 |
|  | Ph.D. | 67 | 19.7 |

Source: Field survey, 2020

### 3.2 Knowledge and awareness on Evidence Informed Decision Making (EIDM)

Table 2 showed the distribution of the respondents on their awareness and knowledge of Evidence Decision Making (EIDM). Approximately 64% of the respondents are not aware of EIDM while 71% had no understanding of EIDM. As regards knowledge of EIDM, 36% had no knowledge of EIDM, 29% had moderate knowledge, 34% were beginners while approximately 1% had advanced knowledge of EIDM. The result showed that the respondents had poor awareness and knowledge of EIDM.

**Table 2:** Knowledge and Awareness on Evidence Informed Decision Making (EIDM)

| Variable | Sub Variable | Frequency | Percentage (%) |
| --- | --- | --- | --- |
| Have you ever heard of EIDM | No | 216 | 63.5 |
|  | Yes | 124 | 36.5 |
| Understand the term EIDM | No | 241 | 70.9 |
|  | Yes | 99 | 29.1 |
| Knowledge of EIDM Rating | Advanced | 2 | 0.6 |
|  | Beginner | 116 | 34.1 |
|  | Moderate | 99 | 29.1 |
|  | None | 123 | 36.2 |

Source: Field survey, 2020

According to Rogers (2003), knowledge occurs when an individual is exposed to an innovation’s existence and gains an understanding of how it functions. The results in this section show that the respondent may not have been exposed to EIDM hence lack/low knowledge of the concept.

Decisions are normally preceded by awareness and gathering of information. When the level of information gathered reaches a significant, the individual’s knowledge on the concept is enhanced leading to correct perception and decisions. Building the respondent’s knowledge on decision making based on informed evidence is paramount to increased policy decisions based on evidence among agencies in the country. Fry, Ryley and Thring (2018) in their study of influence of knowledge and persuasion on the decision to adopt or reject alternative fuel vehicles in Birmingham found that respondent’s poor knowledge and awareness of electric vehicles and their advantages were detrimental to respondents’ adoption of these vehicles.

### 3.3 Perception of EIDM among respondents

Table 3 showed the perception of respondents on EIDM. According to the rating values of the five-point Likert type scale, it was shown that respondents mainly perceived that EIDM should be given more awareness, institutionalized for policy making and implementation and capacity of researchers should be built on EIDM. Ezenweka, *et al*., (2020) in identifying possible solutions to the barriers of EIDM in health sector noted the need for a continuous and sustainable capacity building on EIDM among users and producers of research evidence. In depth knowledge and capacity building of both researchers and users of research evidence in EIDM will help them understand the need and benefit of making decision making based on research evidence. In same vein, Onwujekwe, *et al* (2020) stated that strengthening the capacity of producers and users of research is a more sustainable strategy for developing Health policy and system research in Africa, than relying on training in high-income countries.

**Table 3:** Perception–Respondents perception of EIDM)

| **Perception** | **AGREE** | **DISAGREE** | **Strongly Agree** | **STRONGLY DISAGREE** | **UNDECIDED** | **Rating** |
| --- | --- | --- | --- | --- | --- | --- |
| EIDM actually exists | 194(57.1%) | 15(4.4%) | 3(0.9%) | 5(1.5%) | 123(36.2%) | **3** |
| EIDM is beneficial to policymaking | 269(79.1%) | 28(8.2%) | 1(0.3%) | 4(1.2%) | 38(11.2%) | **4** |
| EIDM could help in formulating good policies | 52(15.3%) | 21(6.2%) | 3(0.9%) | 3(0.9%) | 261(76.8%) | **2** |
| EIDM should be given more awareness | 295(86.8%) | 1(0.3%) | 38(11.2%) | 2(0.6%) | 4(1.2%) | **5** |
| Capacity of policymakers should be built on EIDM | 92(27.1%) | 10(2.9%) | 227(66.8%) | 1(0.3%) | 10(2.9%) | **4** |
| EIDM should be institutionalised for policy making | 297(87.4%) | 1(0.3%) | 29(8.2%) | 1(0.3%) | 12(3.5%) | **5** |
| EIDM should be institutionalised for policy implementation | 308(90.6%) | 3(0.9%) | 25(7.4%) | 1(0.3%) | 3(0.9%) | **5** |
| Capacity of researchers should be built on EIDM | 311(91.5%) | 2(0.6%) | 13(3.8%) | 1(0.3%) | 13(3.8%) | **5** |
| Everybody should know about EIDM | 67(19.7%) | 6(1.8%) | 259(76.2%) | 1(0.3%) | 7(2.1%) | **3** |
| EIDM should be restricted to policy makers | 13(3.8%) | 90(26.5%) | 5(1.5%) | 210(61.8%) | 22(6.5%) | **3** |

Source: Field survey, 2020

### 3.4 Evidence Based Practices among respondents

Result in Table 4 showed respondents’ participation in EIDM practices. Result showed that approximately 99% of the respondents have neither been trained nor involved in training others in EIDM. This buttresses more the importance of capacity building of all parties involved in producing and using research evidence in EIDM. Majority (92%) respondents do not have knowledge of how EIDM have affected their job specification. Earlier results showed that respondents are neither aware nor possess sound knowledge of EIDM and its practices. Therefore, it is not surprising that they may not even be able to identify if it actually affects their job specification.

**Table 4.** Evidence Based practices

| Response on Evidence Based Practices | Variable | Frequency | Percentages |
| --- | --- | --- | --- |
| Have you been trained on EIDM? | No | 338 | 99.4 |
|  | Yes | 2 | 0.6 |
| Have you been involved in training others on EIDM? | No | 338 | 99.4 |
|  | Yes | 2 | 0.6 |
| Have you promoted EIDM before? | No | 89 | 26.2 |
|  | Yes | 251 | 73.6 |
| In what way did EIDM affect your Job specification? | Negatively | 2 | 0.6 |
|  | No knowledge | 314 | 92.4 |
|  | Positively | 20 | 5.9 |
|  | Very negatively | 2 | 0.6 |
|  | Very positively | 2 | 0.6 |
| Institutionalizing EIDM has far reaching implications | No | 299 | 87.9 |
|  | Yes | 34 | 10 |
|  | - | 7 | 2.1 |

Uneke, *et al*., (2016) in their assessment of policymakers’ engagement initiatives to promote Evidence-informed health policymaking in Nigeria found that research on evidence-informed policymaking and knowledge transfer/exchange processes involving policymakers in Nigeria is new or still at infancy stage. This explains the low awareness and knowledge of policy makers in EIDM practices.

### 3.5 Policy-makers’ engagement in EIDM

In this study, it was observed that 8.5% of the respondents were policy-makers in the organization, yet 90.7% of the respondents agree that EIDM is useful in policy-making while 81.3% of the respondents engage their colleagues in EIDM (Table 5). This is an indication that government agencies are moving toward adopting evidence-informed decision making into practice. However, the low numbers of the policy-makers in the organization’s workforce indicates that effort are required to effectively develop capacity and promote contextual factors that advance and sustain EIDM (Dobbins *et al*., 2018). Also, it was observed that over 86% of the respondents do not have access to sources of evidence before making decisions, have not heard or have knowledge on how to synthesize evidence. Similar observations have been reported by Peirson *et al*. (2012), were decision makers in the study often lack access to information, lack the knowledge and skills to conduct systematic literature reviews, and lack infrastructure to support EIDM activities. These are indications of weak linkages that exist among producers (researchers) and users (policymakers) of research evidence which underscores a communication gap in research priority-setting (Ezenwaka *et al*., 2020). Therefore, the level of engagement and interaction between producers and users of research evidence could influence knowledge exchange and sharing, which are critical for bringing about change in evidence-informed policy and practices. Uneke *et al*. (2008) has suggested four strategies for getting research evidence into policy and practice, which are: (i.) increased demand for research evidence by policy-makers, (ii.) involvement of users of evidence in research priority setting, design and implementation, (iii.) facilitating researcher-policy-maker engagement through workshops and research networks, and (iv.) active dissemination of research evidence to relevant policy-makers and other stakeholders.

**Table 5.** Engagement of policy-makers in EIDM

|  |  |  |  |  |  |
| --- | --- | --- | --- | --- | --- |
| Are you a policy maker in your organization/place of work? | NO | YES |  |  |  |
|  | 311(90.7%) | 29(8.5%) |  |  |  |
|  | NO<br>KNOWLEDGE | NOT AT ALL | OFTEN | Very Often | Rating |
| How often do you engage your colleagues in EIDM | 22(6.4%) | 31(9%) | 8(2.3%) | 279(81.3%) | 1 |
| How often do you have access to these sources of evidence? | 33(9.6%) | 295(86%) | 10(2.9%) | 2(0.6%) | 3 |
| Have you heard of evidence synthesis? | 11(3.2%) | 324(94.5%) | 3(0.9%) | 2(0.6%) | 3 |
| How often do you synthesize your evidence | 26(7.6%) | 307(89.5%) | 6(1.7%) | 1(0.3%) | 3 |
|  | NO<br>KNOWLEDGE | NOT USEFUL | USEFUL | Very Useful |  |
| How useful do you see EIDM as a policymaker? | 8(2.3%) | 8(2.3%) | 311(90.7%) | 13(3.8%) | 2 |
|  | DIFFICULT | NO<br>KNOWLEDGE | NOT<br>DIFFICULT | VERY<br>DIFFICULT |  |
| How difficult is it for you to gather enough Evidence before making your decisions? | 300(87.5%) | 29(8.5%) | 4(1.2%) | 7(2%) | 4 |

### 3.6 Policy-makers’ sources of evidence

There is a robust source of evidence used by policy-makers in formulating policies in Nigeria (Uneke *et al*., 2017). Some of the major sources of evidence are shown in Figure 2. From this study, it was observed that all the sources of evidence suggested by the respondents could be categorized into six major groups, namely academic community, media, official statistics, opinion polls, specialized policy unit and traditional knowledge. These groups are similar to the categories of sources of evidence suggested by Gurung (2014). As observed, the major source of evidence is from the academic community made up of the universities and research institutions. However, there is a wide gap between researchers and policy-makers and this has been a major challenge to the development of evidence-based-policy (Uneke *et al*., 2018). Therefore, to promote interactions between researchers, policy-makers and other stakeholders who can influence the uptake of research findings for decision making, the policy-makers should participate in research activities while the researchers get involved in policy-making and implementation processes.

**Figure 2:**
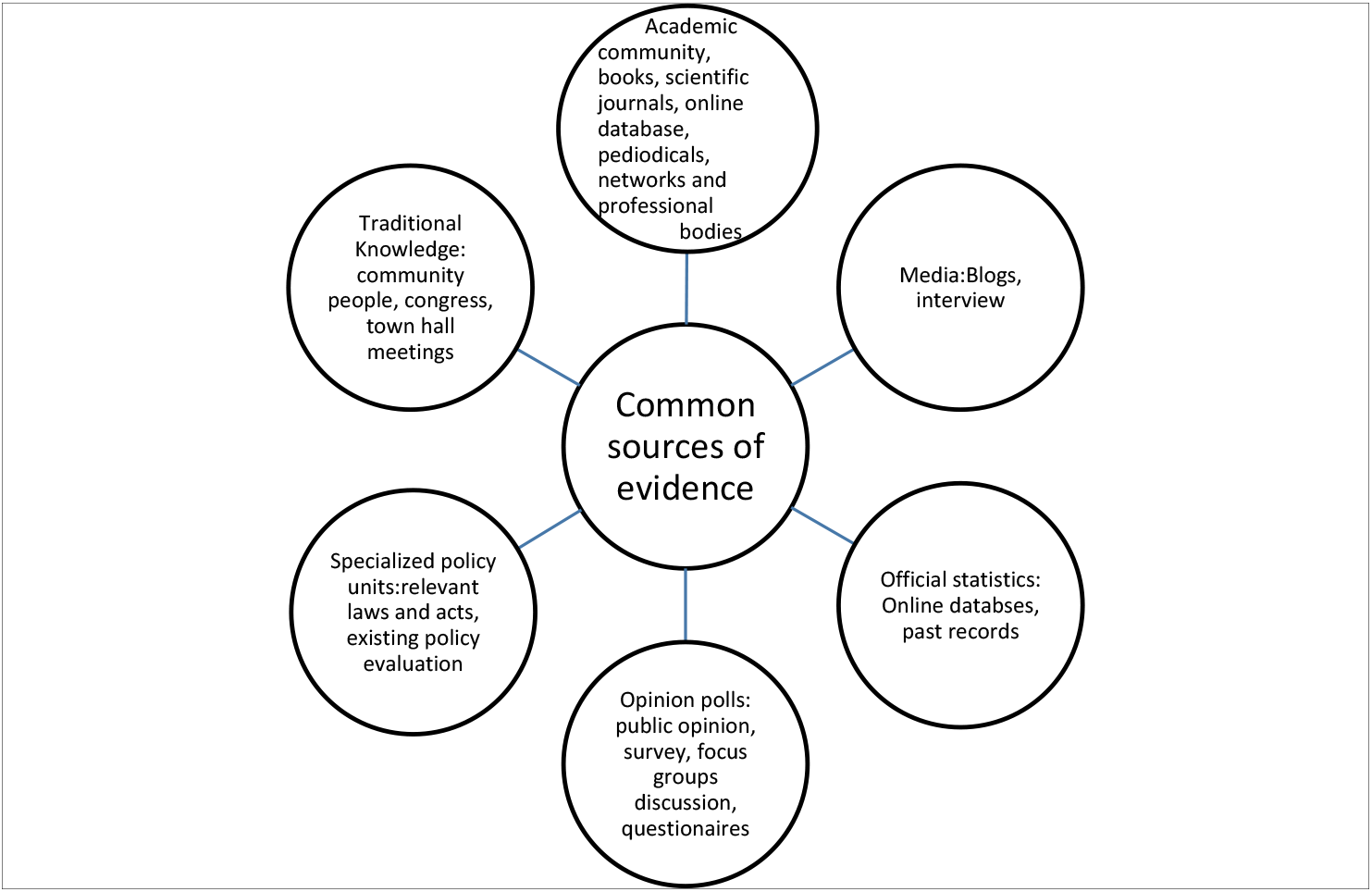
The common sources of research evidence for policy-makers in Nigeria

### 3.7 Institutionalizing EIDM in Government agencies in Nigeria

This study provides further proof of the possible impact of an institutional-wide knowledge translation intervention on the evidence-informed decision making outcomes. It was observed that 91.8% of the respondents (Table 6) were of the opinion that EIDM could make positive grassroots impact on the target audience if there were mass awareness campaigns and sensitization of the public on EIDM. This has been reported to lead to development and strengthening of partnership, capacity and visibility on EIDM in Africa (Motani *et al*., 2019). Although, over 86% of the respondents agreed. that EIDM should be encouraged at every level of policy-making, it was observed that 57.7% of the respondents felt that EIDM delays policy-making. This maybe because of the belief that policymaking is a complicated and lengthy process that is mostly done at national level (Naude *et al*., 2015). However, Peirson *et al*. (2012) and Motani *et al*. (2019) have recommended that for an organization to meet its goal of becoming an evidence informed decision making organization, it will need innovative and long-term capacity-building, dedicated leadership, continuous stakeholder engagement, planning, commitment, and substantial investments.

**Table 6.** Institutionalizing EIDM

|  | AGREE | DISAGREE | NO<br>KNOWLEDGE | STRONGLY<br>AGREE | UNDECIDED |
| --- | --- | --- | --- | --- | --- |
| EIDM is tedious and time wasting | 38(11.1%) | 9(2.6%) | 1(0.3%) | 8(2.4%) | 284(82.8%) |
| EIDM delays policy making | 198(57.7%) | 111(32.4%) | 1(0.3%) | 7(2%) | 23(6.7%) |
| EIDM could make grass root positive impact on the target audience | 18(5.2%) | 2(0.6%) |  | 315(91.8%) | 5(1.5%) |
|  | DISAGREE | Strongly<br>Disagree | UNDECIDED |  |  |
| EIDM should be discouraged | 32(9.3%) | 302(88%) | 6(1.7%) |  |  |
|  | AGREE | DISAGREE | Strongly Agree | STRONGLY<br>DISAGREE | UNDECIDED |
| EIDM should be encouraged at every level of Policy making | 295(86%) | 10(2.9%) | 29(8.5%) | 1(0.3%) | 5(1.5%) |
| EIDM should be institutionalised in all agencies | 75(21.9%) | 38(11.1%) | 221(64.4%) | 1(0.3%) | 5(1.5%) |

Institutionalizing EIDM in government agencies in Nigeria has numerous benefits as shown by the findings of this study. This will only be possible when all parties involved in producing and using research evidence are well informed and knowledgeable in EIDM. Uneke, *et al*., (2011) noted that evidence-based policy is a critical requirement for improved health policy in Nigeria. According to the authors, better use of research evidence in development policy making can save lives through more effective policies that respond to scientific and technological advances, use resources more efficiently and better meet citizens’ needs. One of the most challenging issues associated with institutionalizing EIDM in developing countries like Nigeria is the capacity constraints of policymakers to access, synthesize, adapt and utilize available research evidence (Uneke, *et al*., 2013). According to the authors, a major factor responsible for this is the lack of engagement/involvement of policymakers in the evidence generation process.

Institutionalizing EIDM has far reaching implications and can be practices as follow: -

1. ELA, in each country should meet the national assembly in various countries to make laws that will affect the common man in the society. ELA should become an institution in various countries
2. Enable accurate decision making
3. Enhancement of policy making
4. Every policy of government will be based on results of evidence
5. Facilitates establishment of policy framework
6. help in posting policy
7. helps formulate policy that will have direct impact on the user
8. improves efficiency and competence
9. increased GDP; improved sustainable livelihood for Nigerians; evidence of successful and impactful implementation of public policies; low rate of unemployment rate; inflation
10. it enables the right decision to be made
11. it enhances productivity and competence
12. it has to become part of legislation
13. it will enhance policy making approach in solving problems
14. nation building and good governance
15. no yet specific implications

### 3.8 Strategies for institutionalizing and improving the existence of EIDM

Institutionalization of evidence-informed decision-making (EIDM) involves the integration of the best available research evidence with contextual factors including community preferences, local issues (e.g., health, social), political preferences, and resources (Armstrong *et al*, 2013). Although, there are no set of standard strategies for institutionalizing EIDM in a government agency, however from this study, several approaches were suggested by the respondents and categorized into four steps (Figure 3). The approaches suggested were advocacy and awareness of EIDM, developing legal framework for evidence-informed practices, implementation, and monitoring and evaluation of EIDM. Advocacy and legal framework have been reported to be the most effective means of institutionalization of policies especially in the health sector, because it created a public awareness that was included in the decision-making body (Banken, 2001). Also, in an evidence-informed approach to policy-making, implementation, which refers to the systematic efforts used to encourage adoption of evidence and knowledge by overcoming barriers, is at the core of better decisions for a better future (Stewart, *et al*., 2019). In Nigeria, there are inconsistencies in the implementation of policies amongst the different organs and levels of government in Nigeria, because the implementing agencies in most cases lack appropriate modern technology, managerial skills and administrative capacity that are essential for effective policy implementation, and the procedures adopted in policy implementation are not consistent with policy goals. (Ijie, 2018). However, several ways to implement EIDM were suggested by the respondents. To determine the impact of EIDM, there is a need to monitor and evaluate the progress of the EIDM so as to improve the process. Monitoring policy development and implementation is an integral component of the policy cycle and complements researchers’ ability to link policies with improved service delivery and outcomes that is policy and program evaluation (Health policy project, 2014). Two recommendations were suggested by the respondents.

**Figure 3:**
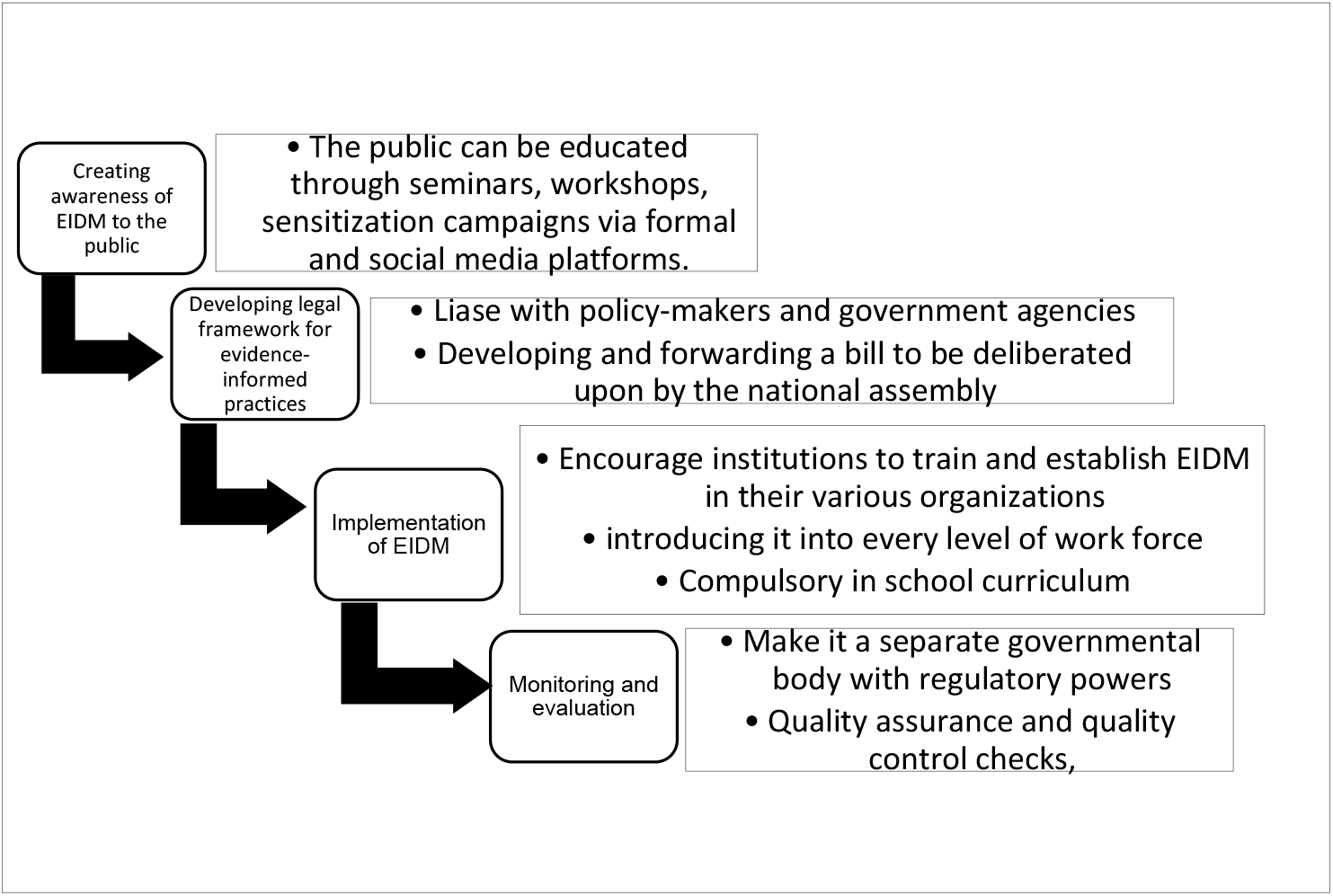
Scheme of the strategies for institutionalizing and improving the existence of EIDM

A close-out conference was held and facilitated sharing of lessons from the researchers’ efforts in translating, promoting, and supporting the use of evidence in government decision-making in their countries. It also featured cementing training skills to further strengthen participating institutions and policymakers as well as researchers on learnt skills to using evidence and sustaining practice.

## 4.0 Recommendations

There is need for a multipronged approach to address the challenges that women face in their quest to pursue STEM courses and succeed while in practice. The study draws out five recommendations that need to be addressed to level the playing field and encourage gender equality in sciences.

- Policy options that target the role of parents and the home environment: provide opportunities for parents to have enough information and social support that they require to assist their children in making decisions on STEM.
- Policy options that target the school environment: fair representation of both gender with reference to the teaching staff of STEM courses. In addition to this, higher education institutions need to award targeted scholarships for women as a booster to the numbers.
- Policy options that target international agencies: these agencies should consider gender equity when awarding grants, especially research grants. The current scenario is gender biased in favour of men which means that less women are likely to grow and compete at higher levels of STEM excellence.
- Workplace policies: proper remuneration of female employees working in STEM is a vital component that needs to be observed by employers for women to be successful in STEM. Employers should also ensure that there is fair representation of women at all levels at work.
- African government policies: African governments need to advocate for and create opportunities for girls to be supported in their pursuit of STEM courses, measures like increasing the budgetary allocation to the education sector will go a long way in increasing the number of women in STEM. There also needs to be deliberate efforts in recruiting women in governance and decision-making positions. Interventions to ensure comprehensive support structures for women in STEM need to be anchored in law through relevant policies to safeguard gender equity in STEM both in the education system and workplaces.

## 5.0 Limitations of Institutionalizing an Evidence Based Practice in the agency

The agency was established in 1997 to enable the Federal Government of Nigeria through Federal Ministry of Science & Technology (FMST) actualize its critical and strategic mandate to research, develop, document. Preserve, conserve and promote Nigeria’s Natural Medicine (Traditional/indigenous Healthcare systems, medications and non-medications healing arts, science & Technology) and assist facilitate their integration into the National Healthcare Delivery System. As well as contribute to the Nation’s wealth and job creation, socio-economic growth and development effort. Evidence-based practice (EBP) is based on scientific evidence which is based on shared vision, decision making between practitioner, patients and others that are significant to them which is based on available information (Enuka and Igbinosun, 2012).

The limitations in institutionalizing EBP in the agency will be discussed under two categories. The first limitations will be discussed under individualized and organizational aspect of the agency. According to previous studies individualized limitations in establishing EBP in the agency has a higher percentage of occurrence compared to the organizational limitations (Khammarnia *et al*., 2015; Abdulwadud *et al*., 2019). Limitations with respect to individualized aspect includes lack of familiarity with EBP, time factors, computer illiteracy, lack of motivation, lack of professional experience and development and negative behavior of staff.

### Lack of familiarity with EBP concept

EBP cannot be established in the agency when the staff and practitioners lack familiarity with the concept of EBP either because they were not trained or they are not just interested in improving their knowledge in EBP. According to Retsas, (2009) stated the term EBP is unfamiliar, it is becoming difficult to integrate evidence successfully. The lack of familiarity with EBP could also be associated to the fact that there are no up to date knowledge of EBP, thus they have difficulty in interpreting their research findings (Enuka and Igbinosun, 2012). There is no updated knowledge of EBP in the agency which can aid researchers to current information which is a barrier to institutionalizing EBP.

### Time factors

A major limitation in establishing EBP in the agency is time constraint. EBP requires a lot of time for reading and research by the staff, researcher and practitioners. Nobody wants to make out time from busy schedules to read relevant articles to get recent information (Khammarnia *et al*., 2015; Abdulwadud *et al*., 2019).

### Computer illiteracy

In an era where the world is going digital and improved technology, some staff and researchers are still computer illiterate. They lack computer based knowledge and skills. If there any staff of the agency has computer constraint they will not be able to access information on electronic version of journals and data information to practice for evidence (Sadeghi-Bazargarri *et al*., 2014; Belizan *et al*., 2007).

### Lack of interest and motivation

Because EBP is not thought in most tertiary institutions in Nigeria makes people not to be interested in learning how to apply evidence. Staff of the agency shows no interest and they are not also motivated by the organization stressing the need and the importance of evidence in research (Ahaive *et al*., 2016; Abdulwadud *et al*., 2017).

### Lack of professional experience

There is lack of professional experience in EBP.

## Organizational Limitations

There are various organizational limitations that can prevent the effective use of EBP in the agency. Such issues that attributed to the organization are: Technology infrastructure, inadequate funding, lack of documentation and workplace constraints (Mehrdad *et al*., 2008; Enuka and Igbinosun, 2012; Khammarnia *et al*., 2015).

### Technology infrastructure

For researchers and practitioners working with the agency to participate in evidence, the technology aspect must be put in place. This includes provision of internet and the necessary information technology in the workplace. The lack of access to information technology that will aid to database and website that contains EBP materials is one of the limitations to the institutionalization of EBP in the agency (Enuka and Igbinosun, 2012).

### Inadequate funding

According to the Director General of the agency Pharm, Sam Oghere in an interview in 2017 spoke extensively on limited funding of the agency. Without sufficient funding it will be difficult to train practitioners and researchers on EBP (Martis *et al*., 2008).

### Lack of documentation

There are no comprehensive inventories of the medicinal plants and natural products in Nigeria. Evidence-based guidelines are readily not available, this has led to the scarcity of information, lack of current journals in the library which has rendered the institutionalization of EBP to no effect. Documentation of EBP can only be achieved when there is coherent government and national policy of the agency in place (Lehane *et al*., 2019).

## Ethics approval and consent to participate

Not applicable

## Consent for publication

Not applicable

## Availability of data and material

The datasets generated during and/or analysed during the current study are not publicly available due to protection of individual privacy but are available from the corresponding author on reasonable request.

## Authors’ contributions

EON and CR conceived the intervention; EON, NF, BO, EOU, JOO, IC, EC, CM and AO designed the work; OEA, PAN and CCI created and validated evaluation documents, CR, CE, VO, IU and AO collected data; EOU, UO, BAS, VO, LUO and CR analyzed and interpreted the data; CCI, EON, LUO, OEA, BAS, BO, JOO, EC, CM, PAN, NF and IC were major contributors in writing the manuscript; All authors participated in systematic reviews of literature read and approved the final manuscript.

## Acknowledgements

The authors are grateful to the technical partners, African Academy of Sciences (AAS) and African Institute for Development Policy (AFIDEP) for phenomenal support and to Dr. Samuel Oghene Etatuvie for necessary approvals.

